# Smooth pursuit eye movements contribute to anticipatory force control during mechanical stopping of moving objects

**DOI:** 10.1101/2023.02.09.527925

**Authors:** Oindrila Sinha, Shirin Madarshahian, Ana Gomez-Granados, Morgan L Paine, Isaac Kurtzer, Tarkeshwar Singh

## Abstract

When stopping a closing door or catching an object, humans process the motion of inertial objects and apply reactive limb force over short period to interact with them. One way in which the visual system processes motion is through extraretinal signals associated with smooth pursuit eye movements (SPEM). We conducted three experiments to investigate how SPEM contribute to anticipatory and reactive hand force modulation when interacting with a virtual object moving in the horizontal plane. We hypothesized that SPEM signals are critical for timing motor responses, anticipatory control of hand force, and task performance. Participants held a robotic manipulandum and attempted to stop an approaching simulated object by applying a force impulse (area under force-time curve) that matched the object’s virtual momentum upon contact. We manipulated the object’s momentum by varying either its virtual mass or its speed under free gaze or constrained gaze conditions. We examined gaze variables, timing of hand motor responses, anticipatory force control, and overall task performance. Our results show that when SPEM were constrained, anticipatory modulation of hand force prior to contact decreased. However, constraining SPEM did not seem to affect the timing of the motor response or the task performance. Together, these results suggest that SPEM may be important for anticipatory control of hand force prior to contact and may also play a critical role in anticipatory stabilization of limb posture when humans interact with moving objects.

**New and Noteworthy:** We show for the first time that smooth pursuit eye movements (SPEM) play a role in modulation of anticipatory control of hand force to stabilize posture against contact forces. SPEM are critical for tracking moving objects, facilitate processing motion of moving objects, and are impacted during aging and in many neurological disorders, such as Alzheimer’s disease and Multiple Sclerosis. These results provide a novel basis to probe how changes in SPEM could contribute to deficient limb motor control in older adults and patients with neurological disorders.

## Introduction

During many activities of daily living, such as stopping a door from closing or catching a ball humans process motion of moving objects and apply force over short time-periods to change their momentum. Success in these scenarios requires that the nervous system estimate object momentum (mass x speed) accurately and use that information to modulate limb force. The visual system has multiple mechanisms to estimate object speed. Humans estimate object speed using retinal cues as the object image moves across the retina. In addition, “extraretinal signals” associated with slow smooth pursuit eye movements (SPEM) that track moving objects also contribute to motion perception. These eye movements could play an important role for motion perception since a reafferent copy of the ocular motor command could provide a predictive signal for motion to be integrated via an internal model (1– 4). In fact, psychophysical studies suggest that extraretinal signals associated with SPEM affect both the perceived speed and direction of object motion (2, 5) and when gaze is experimentally constrained to fixate at a point, then the accuracy of object motion perception is degraded (6, 7).

How humans estimate and compare mass of different objects has also been addressed using psychophysical and neurophysiological studies (8–10). Humans typically associate mass with the size of an object and this contributes to the well-studied size-weight illusion, judging a smaller object as heavier than a larger object of equal weight (11). The neural regions involved in the estimation of mass from its size include the ventral cortex (12) and the frontoparietal areas (13).

During catching, the visuomotor system generates anticipatory commands to counter the reactive forces that would arise during contact (14–16). This involves co-activation of antagonist upper limb muscles (17, 18), where the activity scales with the momentum of the falling object (17), regardless of whether the momentum was manipulated with object speed or mass. This suggests that the visuomotor system estimates the momentum of a moving object before preparing an anticipatory postural response. Anticipatory postural response are a hallmark of motor dexterity and take years to develop (19).

The overall objective of the study was to define how extraretinal signals associated with SPEM contribute to modulate hand force during mechanical interactions with moving objects. Evidence in favor of extraretinal signals associated with SPEM contributing to hand motor responses has come from catching and interception studies (1). Catching studies have shown that participants are less accurate when fixating compared to when they pursue moving targets with SPEM (20, 21). Observers also tend to overestimate target speed during fixation, compared to when eye movements are unrestricted (22–24). This overestimation likely also contributes to participants making interception movements ahead of moving targets when they are fixating instead of pursuing targets (25). Hence, instructing participants to fixate their gaze instead of pursuing will likely alter their anticipatory and reactive responses.

Information may also flow from the hand motor to the ocular motor system. SPEM are substantially improved (fewer catch-up saccades, higher gains, shorter lags between target and gaze) when participants move a target with their own hands compared to when an external agent moves the target (26–29). Accordingly, the ocular motor and hand motor system may reciprocally share information with each other for optimal motion processing and hand motor control.

With this background in mind, our study had two goals. First, determine how the ocular motor system tracks objects moving with different speeds and provides that information to the hand motor system to modulate timing of motor responses and amplitude of anticipatory postural responses. We used anticipatory forces generated by the hand prior to the contact as a measure of anticipatory postural response. An increase in anticipatory grip force prior to contact between the hand and dropped objects has been considered a measure of anticipatory postural control (30–33). Using the idea, we used a robotic manipulandum, which allowed us to measure hand force prior to and during contact with moving virtual objects (see Methods).

Our first hypothesis (H1) was that SPEM modulates the timing of motor responses, anticipatory postural responses, and task performance. Higher SPEM speeds are predicted to modulate the motor timing during free viewing whereas anticipatory responses are predicted to be upregulated with an increase in object mass or speed. When eye movements are constrained, we predict an earlier motor response, and larger anticipatory forces since humans perceive faster object motion during fixed gaze compared to when they pursue an object with SPEM (22–24, 34). The second goal of the study was to probe if manipulating the force requirements during the contact would have an upstream effect on how the ocular motor system tracks moving objects. We hypothesized that selectively lowering the magnitude of force during the contact (while keeping the object momentum and force impulse the same) would lead to lower gaze gains (ratio or gaze and object speed) (H2).

We conducted three experiments to test our hypotheses. In the first experiment (Exp. 1), we examined how SPEM influenced hand posture stabilization. In the second experiment (Exp. 2), we probed how constraining eye movements affected hand posture stabilization. The first two experiments tested the predictions from the first hypothesis (H1). In the final experiment (Exp. 3), we altered the contact dynamics to test upstream effects on eye movements and tested H2.

## Methods

### Participants

36 right-hand dominant participants were recruited to complete the study which consisted of three experiments. Twelve right-handed participants completed the first experiment (23.4 ± 1.5 years; 6 males & 6 females), second experiment (22.8 ± 1.5 years; 6 males & 6 females), and third experiment (24.4 ± 1.5 years; 6 males and 6 females) each. All the participants were screened for any history of neurological conditions or musculoskeletal injuries in the upper limb. Prior to participating, each participant provided written informed consent and was compensated for their time ($10/hr). All study protocols were approved by the Institutional Review Board of the Pennsylvania State University. The participants had to visit the lab twice within 48 hours.

#### Apparatus and stimuli

Participants performed the experimental task (see next section) on a KINARM Endpoint robot (KINARM, Kingston, ON, Canada) with a force sensor embedded in the handle. The KINARM was integrated with an SR EyeLink 1000 Remote eye-tracking system (SR Research, Ottawa, ON, Canada) mounted about 80 cm in front of the participant’s eyes. The Eyelink eye-tracking system is a monocular system with a maximum sampling frequency of 500 Hz, and a precision of 0.5°. Participants grasped a robotic manipulandum with their right hand to interact with virtual stimuli being projected from a monitor on a semi-transparent mirror above the workspace obstructing the direct vision of the hand. Visual stimuli (including the cursor location indicating the current location of the hand) were displayed at 120 Hz. Hand kinematics and kinetics were recorded at 1,000 Hz.

The virtual object presented to the participants was circular and of two different sizes. They were generated using an online Gabor-patch generator (https://www.cogsci.nl/gabor-generator). The orientation of the objects was fixed to 90°, size to 96, Gaussian, standard deviation to 24, frequency to 0.1, and phase to 0. The background color RGB (red, green, and blue) values for the visual object were 0,0 and 0 respectively. The first color was set to 255,0,0 and the second to 255,255,0. The radii of the two objects were set at 1.25 cm and 1.5 cm. The Gabor patch had a clear boundary and was easily detectable in the workspace.

#### Mechanical Stopping of a Virtual Projectile (MSTOP) paradigm

The overall goal of the current study was to understand how smooth pursuit eye movements (SPEM) contribute to hand posture stabilization during interactions with moving objects. To that end, we designed a task called the Mechanical Stopping of a virtual Projectile (MSTOP). Participants were required to stop a virtual object which moved towards them at a constant speed and was assigned an arbitrary mass (see Fig. 1A). The momentum of the object was calculated by multiplying this arbitrary mass and the actual speed of the object. When the object reached the hand, the momentum of the object was converted into a force impulse (using Eq. 1a) that the robot applied on the hand in the form of a trapezoidal contact force over a fixed time duration (90 ms in Exps 1 & 2 and 160 ms in Exp. 3, 10 ms rise and fall times, red curve in Fig. 1A). This simulated a contact between the object and the hand. Participants were instructed to apply a reactive force on the manipulandum (blue curve, Fig. 1A) to match this impulse (Eq. 1b) within an ±8% error margin (see inset Fig. 1A & Eq. 1c). This margin was chosen based on pilot studies that showed that the participants could learn the task at a reasonable rate during a single session of 1.5-2 hours.

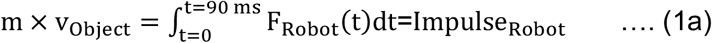

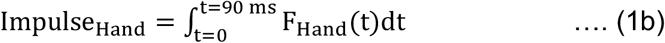

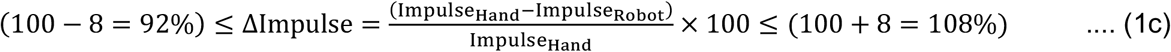

**Figure 1:**
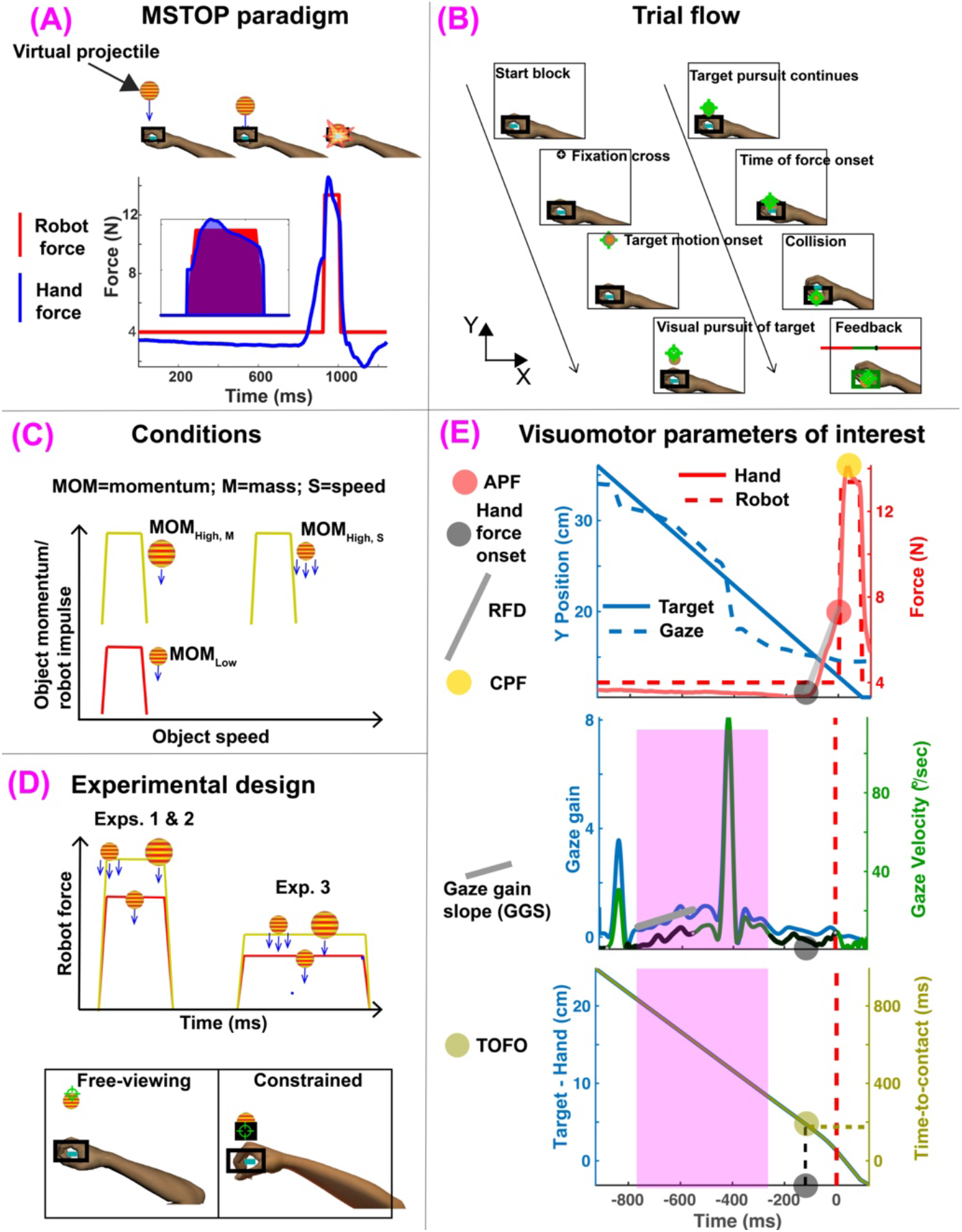
Experimental framework. A) MSTOP paradigm. Participants were required to stop a virtual target moving towards them. When the target reached the hand, the robot applied a force impulse on the hand based on Eq. 1. Participants were required to apply a force on the manipulandum (blue curve) to match the robot impulse (area under the red curve). B) Trial Flow. At the beginning of each trial, participants were instructed to move a cursor representing the veridical position of the hand (blue rectangle) to a start position (black outlined rectangle). Then a fixation cross appeared. Participants had to foveate the cross. After the cross disappeared, the target came on and moved towards the participant. C) Experimental Conditions. The three momentum conditions in all three experiments were created by varying the momentum of the moving target by manipulating either its virtual mass or actual speed. We had a low momentum condition (MOM_Low_) shown in red (served as the baseline condition) and two high momentum conditions (MOM_High,M_ with large target mass and size, and MOM_High,S_ with high speed, both are shown in yellow). Note that these two high momentum conditions had the same momentum. D) Experimental design. The study had three experiments. In Experiments 1 and 2, the contact between the hand and the target occurred over 90ms. Day 2 of Exp. 2 was different from Exp. 1 as it had four blocks of free-viewing, eight blocks of constrained gaze, and then again four blocks of free-viewing (also see Table 1). Exps. 2 and 3 had the same experimental conditions and design, but the contact was softer in Exp. 3 and occurred over 160ms. E) Visuomotor parameters of interest. Top panel shows the target and gaze Y position on the left y-axis and the robot and hand force on the right y-axis for one trial. Rate of force development (RFD), anticipatory peak force (APF), and peak reactive force during contact (CPF) are shown. Middle panel shows gaze gain on the left y-axis and gaze velocity on the right y-axis. Gaze gain slope (GGS) is shown in grey. The pink window indicates the duration over which SPEM signals were considered for analyses. Bottom panel shows how time of force onset (TOFO) was calculated based on the distance between the target and the hand (left y-axis) at the time of hand force onset (HFO) and the time-to-contact between the target and the hand (right y-axis).

At the beginning of each trial, participants were instructed to move a cursor (blue rectangle (see Fig. 1B), 3×1.5 cm) representing their veridical hand position to a start position (black outlined rectangle, 10×6 cm) located at the midline of the visual display (X=0, Y=0). After the start position was reached, a background load (4 N in the -Y direction) was applied and stayed on for the remainder of the trial.

Background loads of 4N were applied to stabilize the hand prior to the presentation of the moving object (35–37) and to minimize unnecessary anticipatory hand movements that would affect the measurement of force onset. Then a fixation cross appeared along the midline 19 cm from the start position further away from the body. Participants were required to fixate their gaze on the cross for 800 milliseconds (ms) and while their hand stayed in the rectangle. 250 ms after the fixation cross disappeared, the virtual object generated using the Gabor-patch generator appeared on the KINARM workspace (18cm) from the hand of the participant and started moving vertically towards the start box. Participants were instructed to “match the handle’s force to stop the ball”. If ΔImpulse was within 92-108%, the trial was successful, and the object stopped. If it exceeded 108%, the object bounced back. If it was less than 92%, the object continued unabated on its original trajectory. Participants were also provided quantitative feedback after every trial in the form of an error scale (see Fig. 1B bottom right).

#### Experimental conditions

We had three conditions in each experiment where we varied the object momentum by either varying its speed or mass. MOM_Low_ was the baseline condition where the object’s virtual mass was fixed at 3 kg and its speed at 25 cm/s. The radius of this object was 1.25 cm. MOM_High, M_ was the first of the two high momentum conditions where the object mass was increased to 4.2 kg, but the speed was the same as MOM_Low_, 25 cm/s. The larger object mass had a larger radius (1.5 cm). The size of the object was increased to reflect its higher mass. MOM_High, S_ was the final high momentum condition where the object mass was the same as MOM_Low_, 3 kg, but its speed was higher at 35 cm/s (see Fig. 1C).

#### Study design

Exps. 1 & 2 were designed to address the first hypothesis (H1). Exp. 3 was designed to test H2 (see top panel, Fig. 1D). Each experiment was conducted over two days. The first day was identical for all experiments. Briefly, participants performed 8 blocks (24 trials each) of MOM_Low_ first and then performed 4 blocks each of MOM_High,S_ and MOM_High, M_ in a randomized order (see Table 1). Day 1 was necessary to get participants trained on this novel task. All participants tracked the objects with smooth pursuit eye movements (SPEM). We called this the free-viewing (FV) condition (Fig. 1D).

On Day 2 of Exp. 1, participants performed 20 blocks of trials where each block consisted of 16 trials of MOM_Low_, and 4 trials each of MOM_High,S_ and MOM_High, M_ presented in a pseudorandom order within a block. This was under free viewing.

Day 2 for Exps. 2 and 3 had identical designs. Participants first performed four blocks of free-viewing (FV1) trials where each block consisted of 16 trials of MOM_Low_, and 4 trials each of MOM_High,S_ and MOM_High, M_ presented in a pseudorandom order. Then in the next 8 blocks we asked participants to fixate at a specific location during the trials. We called these blocks constrained (CONST) viewing blocks. The fixation cross appeared 5 cm from the black rectangle in the Y direction as shown in the bottom panel of Figure 1D. Participants were instructed to fixate their gaze at the cross throughout the trial. A black rectangle (width was double the diameter of the virtual object and the length was 8cm) was placed underneath the fixation cross which would act as an occluder when the object passed the fixation cross. The location of the occluder ensured that smooth pursuit eye movements (SPEM) were minimized in the CONST conditions. The occluder was also strategically placed to ensure that the object image moved on the fovea just before the contact. This sometimes also triggered very brief SPEM, but the durations were much smaller compared to the free-viewing conditions (see Fig. 6, Results). This form of fixation has been used in previous studies to suppress eye movements during motion-perception tasks (24, 38). During the CONST blocks, the extraretinal signals associated with SPEM were minimized, and participants could only judge motion through retinal slip of the object’s image on the retina (retinal motion). At the conclusion of the 8 blocks, the participants performed 4 blocks of free-viewing (FV2) trials again.

### Data recording and analysis

Hand variables were categorized as either anticipatory (or preparatory) or reactive variable(s). Anticipatory variables consisted of both kinematic and kinetic variables. For pre-processing, kinetic variables were low-pass filtered at 50 Hz and kinematic signals were low-pass filtered at 15 Hz. The data were all double-pass filtered using a 3rd order filter with zero-lag filtering. In anticipation of the contact, participants monotonically increased the hand force above the baseline -4 N load and made a small movement towards the object. This force almost always peaked right before the contact between the object and the hand. The hand force at the instance of contact was called the anticipatory peak force (APF) (see top panel, Fig. 1E). We then traced the force signal backward in time and found the first time point when the hand force dropped below 5% of the peak force value (5% of APF). Then we found the nearest minimum force by finding an inflection point in the force-time signal. The minimum of 5% of APF and the inflection point was called the time of hand force onset (HFO). We fit a first-order polynomial to the force data between HFO and APF and computed the slope of the fitted line. This slope provided a measure of rate of force development (RFD) between preparatory HFO and APF. Peak hand reactive force during contact (CPF) was measured as the largest reactive force during the contact. RFD and APF were the two kinetic anticipatory variables of interest. CPF was the only kinetic variable of interest during the contact. A summary of how these variables were calculated is shown in Table 2.

**Table 1:** Overall study design with three experiments conducted over two days.

| **Exp. 1** |  | (The order of these two sets of blocks was randomized) |  |
| --- | --- | --- | --- |
| Day 1 | 8 blocks of MOM<sub>LOW</sub> (24 trials), free viewing | 4 blocks of MOM<sub>High,S</sub> (24 trials), free viewing | 4 blocks of MOM<sub>High,M</sub> (24 trials), free viewing |
| Day 2 | 20 blocks of randomized free-viewing (16 trials of MOM<sub>LOW</sub>, 4 trials MOM<sub>High,S</sub>, 4 trials of MOM<sub>High,M</sub>) |  |  |
| **Exp. 2** |  | (The order of these two sets of blocks was randomized) |  |
| Day 1 | 8 blocks of MOM<sub>LOW</sub> (24 trials), free viewing | 4 blocks of MOM<sub>High,S</sub> (24 trials), free viewing | 4 blocks of MOM<sub>High,M</sub> (24 trials), free viewing |
| Day 2 | 4 blocks randomized free viewing | 8 blocks of randomized constrained | 4 blocks of randomized free viewing |
| **Exp. 3** |  | (The order of these two sets of blocks was randomized) |  |
| Day 1 | 8 blocks of MOM<sub>LOW</sub> (24 trials), free viewing | 4 blocks of MOM<sub>High,S</sub> (24 trials), free viewing | 4 blocks of MOM<sub>High,M</sub> (24 trials), free viewing |
| Day 2 | 4 blocks randomized free viewing | 8 blocks of randomized constrained | 4 blocks randomized free viewing |

**Table 2:** List of all variables and how they were calculated.

| **Variable** | **Calculation** |
| --- | --- |
| GGS | Calculated from slope of gaze gain (gaze speed/object speed) in a time window between 150 ms after object onset and 150 ms before HFO |
| PP | The overall proportion of time the gaze pursued the object |
| HFO | The time at which hand force increased above the 4N baseline values. Only calculated to obtain TOFO and RFD. HFO is not analyzed in the paper |
| TOFO | Distance between object and hand at HFO divided by object speed |
| RFD | Linear polynomial fit between the hand force value at HFO and APF |
| APF | Hand force just prior to impact between object and hand |
| CPF | Peak hand force during contact (90 ms for Exps. 1 & 2 and 160 ms for Exp. 3) |

At the time of hand force onset (HFO), we calculated the distance between the hand and the object along the y-direction (see Fig. 1B) and divided it by the object’s speed to obtain the time of force onset (TOFO, see bottom panel, Fig. 1E). Therefore, this time was a measure of the time-to-contact between the object and the hand at the time point at which participants increased hand force. Increasing the hand force over the baseline force also produced hand motion towards the object. We calculated hand motion onset using the same criterion as HFO and found that those two times were almost perfectly correlated (0.96). Therefore, we chose TOFO as the only hand kinematic variable of interest.

The gaze data preprocessing, transformation and classification were done as described in our previous publications (35, 39). Briefly, gaze point-of-regard (POR) data were low-pass filtered at 15 Hz. We then converted the Cartesian gaze POR data to spherical coordinates and inverse computed the ocular kinematics. The angular gaze speed was then filtered using the Savitzky Golay 6^th^ order filter with window size (or frame length) of 27. We then identified different gaze events (saccades, fixations, smooth pursuits) by using a thresholding technique (39). Within a single trial participants made both smooth pursuit eye movements and saccades and fixated on either the fixation cross or the cursor that represented the veridical location of the hand (see middle panel, Fig. 1E). Our algorithm robustly identified these different gaze events. For smooth pursuit eye movements (SPEM), we calculated gaze gain (ratio of gaze speed and object speed in the Y direction) at each time point. To eliminate confounds associated with using a remote eye-tracker which allowed small head movements to occur, we report gaze gain instead of SPEM gain as has been done by us and others (35, 40).

Since the object was moving in the transverse plane towards the participant, gaze gain was very likely to change as the object got closer to the body. Thus, to obtain a measure of how gaze gain was modulated during the trial, we quantified the slope of the gaze gain. As is often the case, SPEM were disrupted by catch-up saccades (41, 42), especially when the object moved at faster speeds during the MOM_High, S_ conditions. Since visual processing is minimized during saccades, we calculated gaze gain slope (GGS) for only those trials where at least 50% of the gaze event were classified as SPEM by our algorithm. GGS (middle panel, Fig. 1E) was calculated by fitting a linear regression of the gaze gain in a time interval 150 ms after the object appeared and 150 ms before the participants increased their hand force in anticipation of contact between the object and the hand (pink shaded area in middle panel of Fig. 1E). The 150 ms windows were chosen based on published visuomotor reaction times (reviewed in (43)). Within this time window, SPEM were disrupted by catch-up saccades. We calculated a GGS for each instance when the gaze was in smooth pursuit and then used a weighted average to calculate a net GGS for this time window as shown in equation 2.

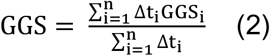

Here Δ t_i_ is the time duration of a continuous smooth pursuit during the above-mentioned time window and n was the total no of pursuits interspersed by catch-up saccades. Each Δ t_i_ had to be at least 40 ms for the pursuit to be included into the calculation.

### Hand force control model

Efferent copies of extraretinal signals associated with smooth pursuit eye movements (SPEM) may serve as feedforward signals to the hand motor system for action control (1, 44). Here, we propose that these extraretinal signals may contribute to hand motor control through the gaze gain signal, which may contribute to the timing of the motor response (TOFO, see Fig. 2A). We further predicted that TOFO would directly predict the rate at which participants increase hand force (RFD) and the hand force right before contact (APF). Stabilization of hand posture in anticipation of contact between the hand and a moving object is critical (31, 32). We expected that participants would increase hand force prior to contact to stabilize posture. Though RFD and APF have no actual bearing on task performance (see Eq. 1), they likely serve a critical need for posture stabilization. Indeed, every single participant increased hand force prior to contact. We predicted that APF, which would measure anticipatory posture stabilization, would also predict ΔImpulse (see Eq. 1c), i.e., how successful participants were at the task (right panel, Fig. 2A).

**Figure 2:**
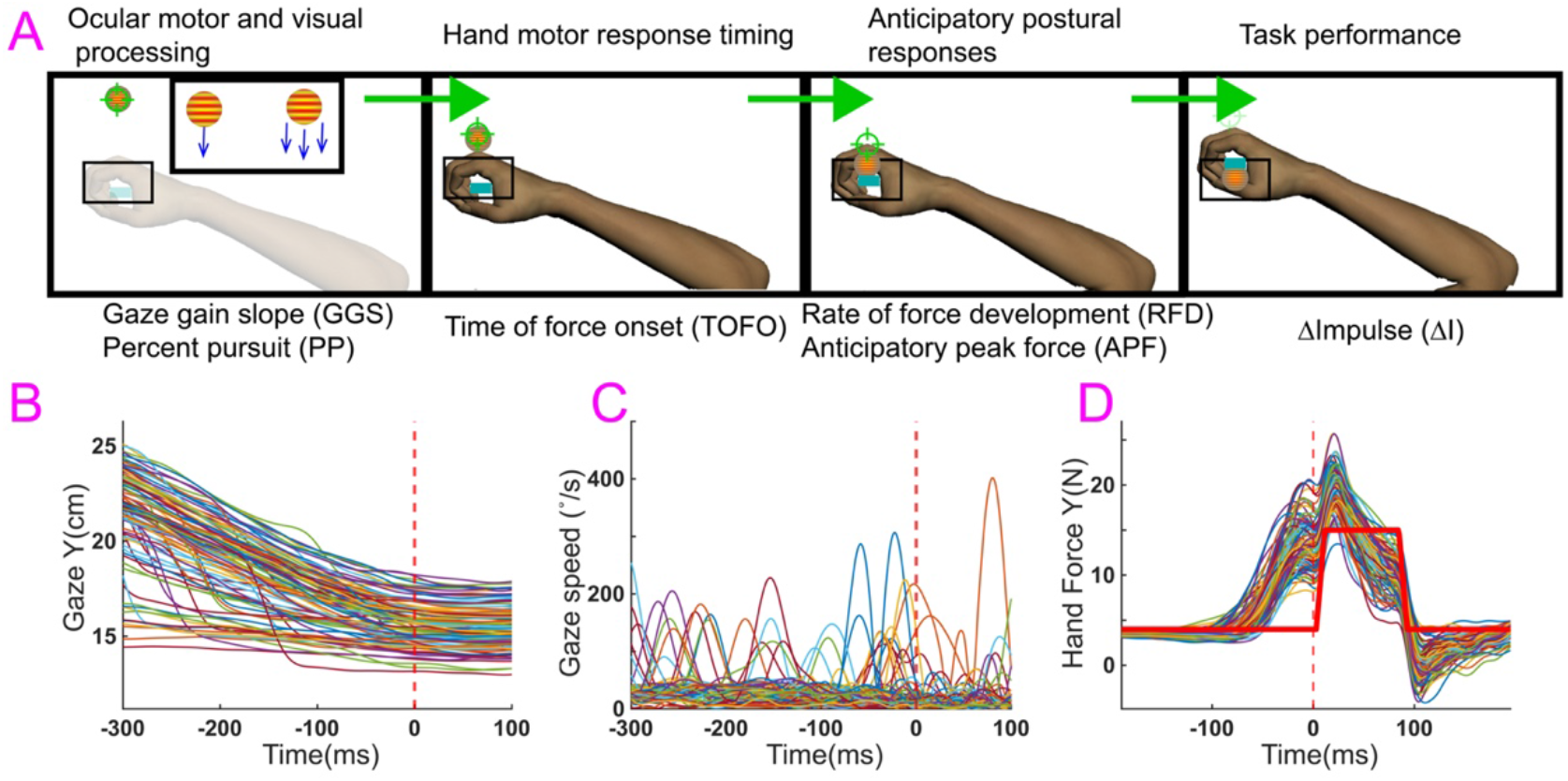
Proposed visuomotor force control model and exemplar trials. A) Gaze gain slope (GGS) and the percentage of time the gaze pursued the target (PP) would determine the time at which participants increase hand force above baseline levels (TOFO) in preparation of the contact between the hand and the target. This time then would determine the rate at which the hand force (RFD) increases and the peak hand force just prior to contact (APF) between the hand and the target. RFD and APF are measures of how the nervous system stabilizes posture in preparation of the contact. Finally, ΔImpulse, the percentage difference between the force impulse applied by the robot and the reactive impulse applied by the hand determines whether the task is successfully performed. B, C, and D, Example trials showing the gaze position, gaze speed, and hand force in the y direction of one block of MOM_High,M_ condition.

#### Statistics

Day 1 and Day 2 for all the experiments were analyzed separately. Day 1 analysis only focused on determining how participants learned the task. To that end, we performed a one-way mixed model repeated measures ANOVA between the first and the last block with *blocks* as the fixed effect and *participants* as random effects, with *participants* nested inside *blocks*.

For Day 2 of Exp.1, we used hierarchical bootstrap sampling with replacement (45, 46) to correlate dependent variables in our visuomotor force control model (Fig. 2A). We also performed a one-way mixed model repeated measures ANOVA with *momentum* (MOM_Low_, MOM_High,M_, and MOM_High,S_ as levels) as the fixed effect and participants as random effects with participants nested inside *momentum*. The repeated measures ANOVA compared the effect of modulating object momentum by changing its mass and speed on our dependent variables.

The existence of moderate to strong correlations between our dependent variables is a necessary but not a sufficient condition to demonstrate the viability of the hand force control model (47). To that end, we constrained gaze in Day 2 of Exp. 2 in different blocks and looked at how the relationship between the variables in the visuomotor force control model changed. We performed a two-way mixed model repeated measures ANOVA with *momentum* (MOM_Low_, MOM_High,M_, and MOM_High,S_ as levels) as one fixed effect factor, *gaze condition* (free-viewing and constrained viewing) as the second fixed effect factor, and participants as random effects. The RM ANOVA compared the effect of manipulating object momentum by changing its mass (MOM_Low_ to MOM_High,M_) and speed (MOM_Low_ to MOM_High,S_), and the effect of constraining SPEM on our variables of interest.

Finally, because it has been hypothesized that the ocular motor and hand motor systems may reciprocally exchange motor information, we posited that changing the reactive force during contact would have upstream effects on eye movements and anticipatory postural responses. We compared the dependent variables in Exps. 2 and 3. We used a one-way ANOVA test in R as well as the non-parametric two-sample Kolmogorov-Smirnov test using the kstest2 function in MATLAB. This test provides a decision for the null hypothesis whether two groups of data are from the same continuous distribution. The one-way ANOVA test in R utilizes un-pooled variances and a correction to the degrees of freedom which is required when equal variances cannot be assumed as was the case when we compared data from Exps. 2 and 3.

The alpha level was set at 0.05, and effect sizes are reported using generalized η^2^. Shapiro-Wilk’s test and Levene’s test were used for checking normality and homoscedasticity assumptions, respectively. Any deviation from the assumptions was corrected using transformations. We used percentile bootstrap, a non-parametric method based on simulations to estimate 95% confidence intervals for correlation coefficients (45, 46). This method allows us to make inferences about a parameter using experimental data, without making assumptions about underlying distributions and works well for making inferences about correlation and regression coefficients (45). Post-hoc pairwise comparisons were conducted using a pairwise t-test to determine the difference between the momentum conditions and the Bonferroni adjusted p-values for multiple comparisons are reported in the result.

For each momentum condition (M_Low_, M_High,M_ and M_High,S_), we pooled all the trials from all the blocks and calculated the Pearson correlation coefficient between pairs of variables of interest (e.g., GGS and TOFO) for each participant. Thus, we get 12 correlation coefficients, one for each participant. Next, we sampled 12 coefficients with replacement and repeated and calculated the median correlation coefficient. We observed that our confidence intervals robustly converged with 2000 repetitions, so we repeated the process 2000 times. We create a 95% non-parametric confidence interval of these 2000 median coefficients. If the CI included ‘0’, we determined that the correlation between the variables was not significant and if it did not include ‘0’, we determined that it was statistically significant.

We performed a regression on ΔImpulse with APF and CPF as predictor variables for each momentum condition for each participant.

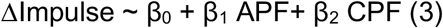

We repeated the bootstrap process 2000 times on β_1_ and β_2_ and calculated a 95% CI. If the CI included ‘0’, we determined that the predictor variable was not a significant predictor of ΔImpulse.

## Results

The goal of the study was to define how smooth pursuit eye movements (SPEM) contribute to timing of motor responses, anticipatory hand force control prior to contact with a moving object, and overall task performance. Figure 2 shows exemplar gaze position (Fig. 2B), gaze angular speed (Fig. 2C) and hand force data (Fig. 2D) of a single participant during Day 1 of Exp.1 in the M_High,M_ condition.

### Day 1 performance shows that participants learned the task by reducing task performance variability

The success rate for Exp. 1 (Fig. 3A) showed a main effect of the first and the last block for MOM_Low_ (*F*(1,11) = 107.44, *p* < 0.001, η^2^ = 0.65) and MOM_High,S_ (*F*(1,11) = 8.65, *p* < 0.05, η^2^ = 0.19), but not for MOM_High,M_ (*F*(1,11) = 4.08, *p* =0.07, η^2^ = 0.09). We found similar patterns of improvement in success rate for Exp. 2. There was a main effect of blocks for MOM_Low_ (*F*(1,11) = 12.71, *p* < 0.05, η^2^ = 0.24) and MOM_High,S_ *F*(1,11) = 19.31, *p* < 0.05, η^2^ = 0.26), but not MOM_High,M_ (*F*(1,11) = 0.9, *p* =0.36, η^2^ = 0.02). For Exp. 3, we also found a main effect of blocks for MOM_Low_ (*F*(1,11) = 36.41, *p* < 0.001, η^2^ = 0.35) and MOM_High,S_ *F*(1,11) = 9.8, *p* < 0.05, η^2^ = 0.15), but not for MOM_High,M_ (*F*(1,11) = 3.6, *p* = 0.08, η^2^ = 0.04).

**Figure 3:**
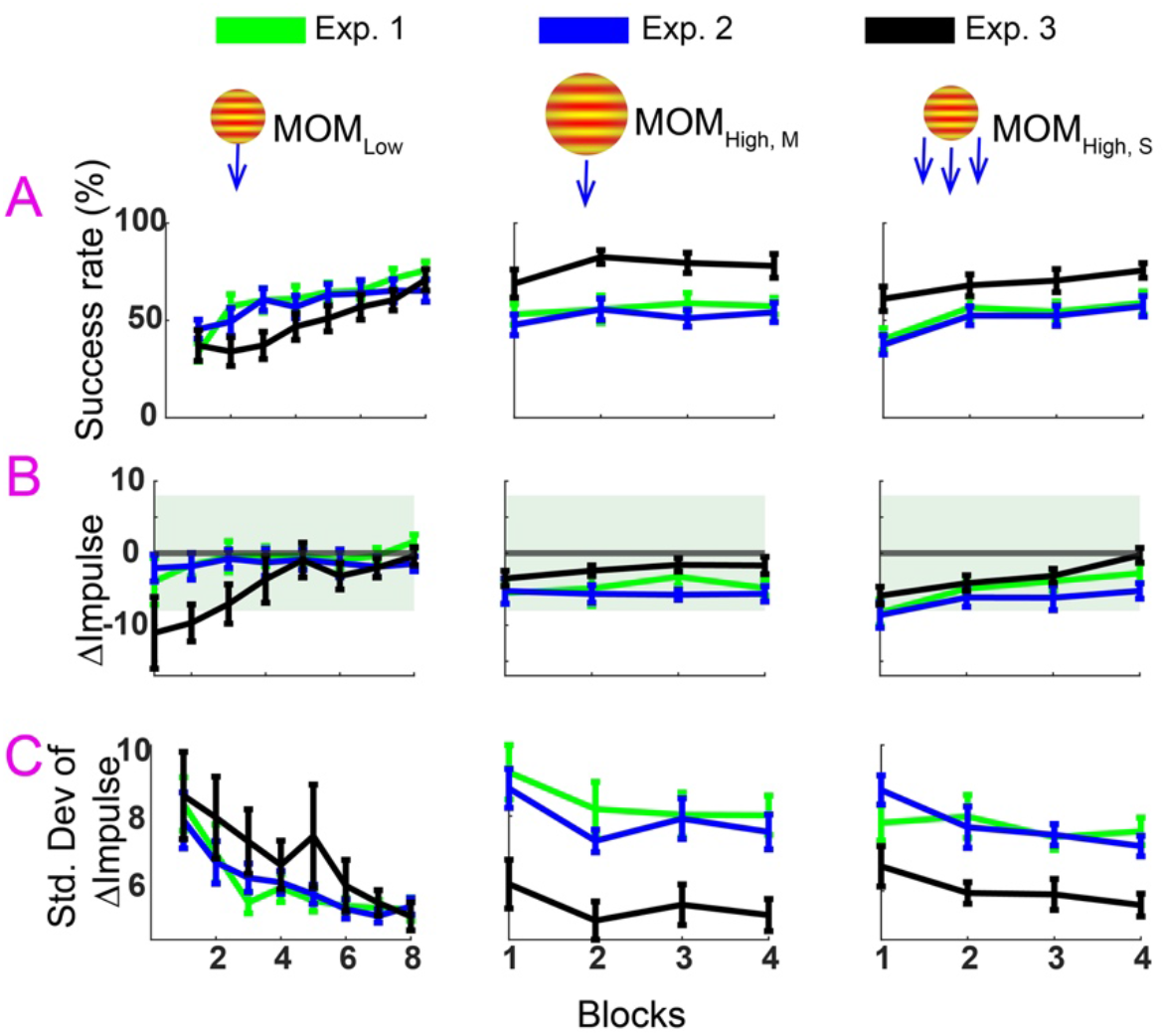
Day one performance showed that participants learned the task by minimizing variability in task performance. A) Success rate in blocks improved with practice for MOM_Low_, but not for the two high momentum conditions (MOM_High,M_ and MOM_High,S_). B) ΔImpulse did not improve with practice for any of the momentum conditions in all three experiments. C) The standard deviation of ΔImpulse within a block showed the most consistent change across all three experiments and momentum conditions.

The same analysis on ΔImpulse (Fig. 3B) showed no main effects of blocks. This suggested that participants may have improved their success rate by minimizing the variance of ΔImpulse across blocks rather than the mean magnitude. Thus, we calculated the standard deviation of ΔImpulse for each block and compared that across blocks (Fig. 3C). The standard deviation of ΔImpulse for Exp. 1 showed a main effect of blocks for MOM_Low_ (*F*(1,11) = 26.49, *p* < 0.001, η^2^ = 0.37), but not for MOM_High,S_ (*F*(1,11) = 0.18, *p* =0.68, η^2^ = 0.007) or MOM_High,M_(*F*(1,11) = 2.39, *p* = 0.15, η^2^ = 0.07). In Exp. 2, there was a main effect of blocks for MOM_Low_ (*F*(1,11) = 11.94, *p* < 0.05, η^2^ = 0.28), MOM_High,S_ (*F*(1,11) = 11.54, *p* < 0.05, η^2^ = 0.31) and MOM_High,M_ (*F*(1,11) = 7.42, *p* < 0.05, η^2^ = 0.11). Similarly for Exp. 3, we found a main effect of blocks for MOM_Low_ (*F*(1,11) = 11.63 *p* < 0.05, η^2^ = 0.24), MOM_High,S_ (*F*(1,11) = 6.01, *p* < 0.05, η^2^ = 0.11), as well as MOM_High,M_ (*F*(1,11) = 5.13, *p* < 0.05, η^2^ = 0.05).

#### Moderate to strong correlations in Exp. 1 supported the hand force control model

Linear correlations and percentile bootstrap analyses support the hand force control model. The first component of the hand force control model predicts a linear relationship between gaze gain slope (GGS) and the time of force onset (TOFO). We correlated these two variables for all trials for each participant. The left panel of Figure 4A shows the Pearson correlation between GGS and TOFO for each participant for the M_High,M_ condition. The bootstrapped confidence interval for the correlation for all 12 participants is shown in the right panel of Figure 4A. The non-parametric 95% confidence interval did not include ‘0’ consistent with a significant correlation. Similarly, we checked the relationship between TOFO and rate of force development (RFD) (Fig. 4B), RFD and anticipatory peak force (APF) (Fig. 4C), APF and ΔImpulse (Fig. 4D) and found these relationships to be significant as well. TOFO and RFD were negatively correlated, while RFD and APF, and APF and ΔImpulse were positively correlated. Table 3 shows the bootstrap intervals and median correlations for all sets of variables for all three conditions (MOM_Low_, MOM_High,M_, and MOM_High,S_).

**Figure 4:**
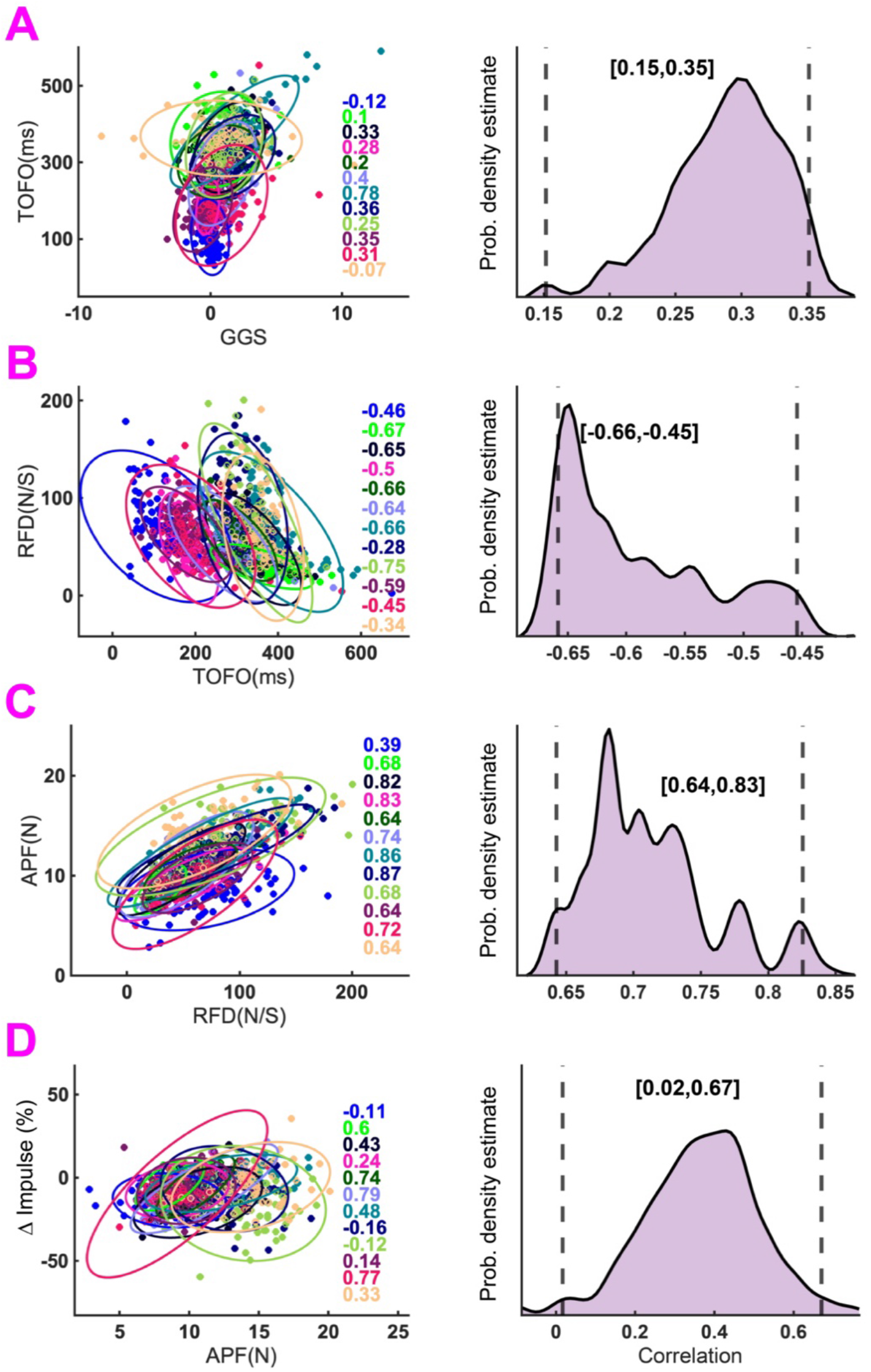
Correlations supported the hand force control model. Left panels show the Pearson correlation between variables of interest for all 12 participants in Exp.1, Day 2, M_High,M_ condition. Each dot represents a trial, and each color represents a participant. The right column shows the bootstrapped distribution of the individual correlation with non-parametric 95% confidence intervals (CI) in brackets. We consider that there was a significant correlation between the variables if the CI did not include 0. A) There was a significant and moderately strong positive correlation between gaze gain slope (GGS) and time of force onset (TOFO). B) TOFO and rate of force development (RFD) were negatively correlated – when TOFO was short, participants increase hand force more rapidly. C) RFD and anticipatory peak force (APF) were also strongly and positively correlated. D) The correlation between APF and ΔImpulse (task performance) was positive and significant, but weak. The correlations for the other two momentum conditions in Exp. 1 are shown in Table 1.

**Table 3:**
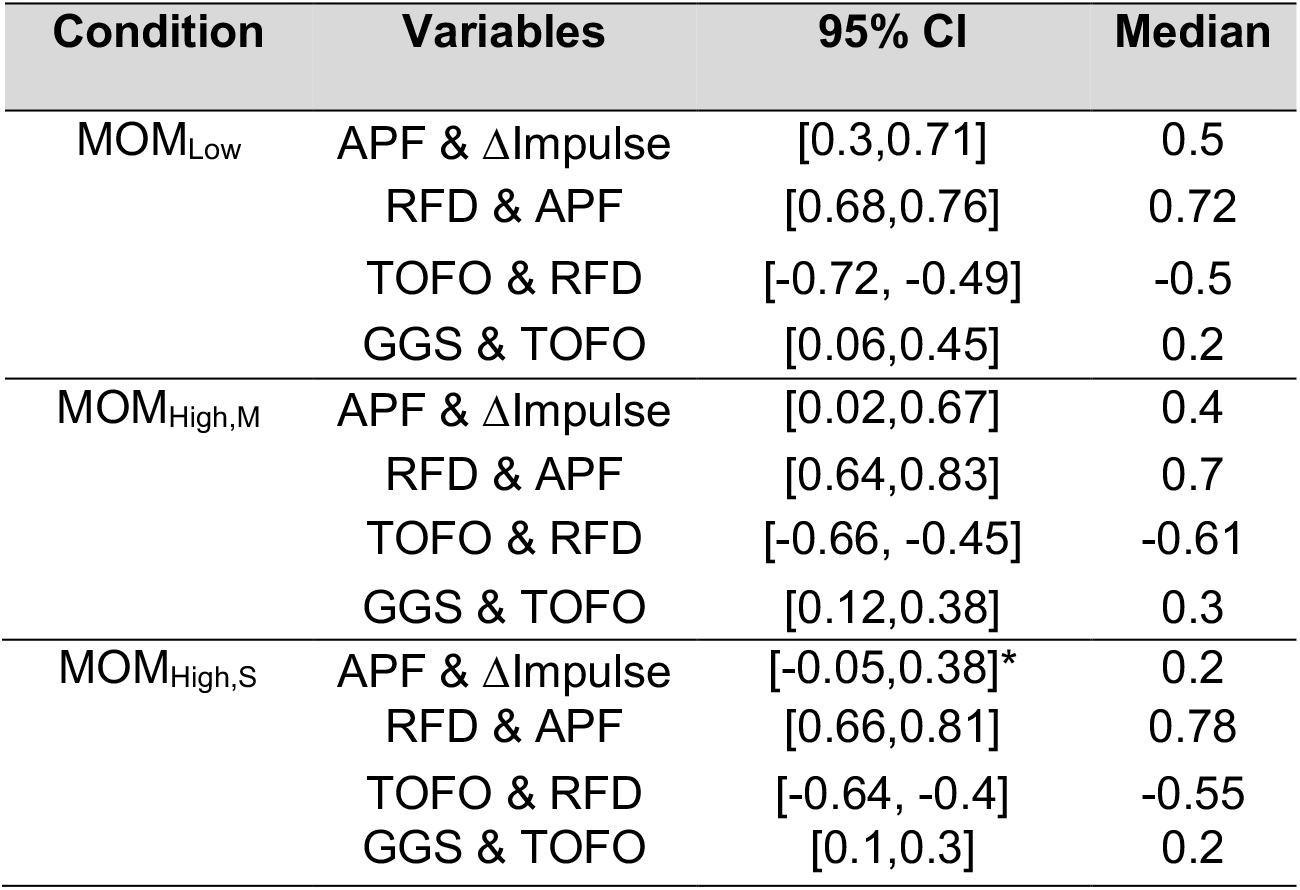
Bootstrap confidence intervals for all correlations and all conditions. * The only interval that spanned ‘0’

The correlations between TOFO and RFD, and RFD and APF are the strongest in all three conditions whereas the correlation between GGS and TOFO is weaker. These results support the hand force control model and suggest that smooth pursuit eye movements may contribute to the modulation of anticipatory postural responses (RFD and APF) and task performance (ΔImpulse).

### Object speed affected timing of the motor response while object momentum affected the anticipatory postural responses in Exp. 1

When object momentum was increased by a speed increase, the slope of the gaze gain (GGS) increased (upper panels of Fig. 5A) [main effect of momentum for GGS: *F* (2,22) = 50.3, *P* < 0.001, η^2^ = 0.64], while the time of force onset (TOFO) decreased (Fig. 5A) [main effect of momentum for TOFO: *F*(2,22) = 13.18, *P* < 0.001, η^2^ = 0.05]. Post hoc pairwise t-test showed a significant difference between (MOM_High,S &_ MOM_Low_, and MOM_High,S &_ MOM_High,M_ for both GGS (p<0.001) and TOFO (p<0.01). In contrast, anticipatory control variables (RFD and APF) increased in magnitude with the momentum of the object (lower panels of Fig. 5A), irrespective of whether the momentum increased due to speed (MOM_High,S_) or mass (MOM_High,M_). The one-way RM ANOVA results showed a main effect of momentum for RFD (*F* (2,22) = 19.64, *P* < 0.001, η^2^ = 0.41) and main effect of momentum for APF (*F* (2,22) = 31.4, *P* < 0.001, η^2^ = 0.40). Post-hoc comparisons revealed a significant difference between MOM_Low_ and MOM_High,M_ and MOM_Low_ and MOM_High,S_ for both APF and RFD (p<0.01).

**Figure 5:**
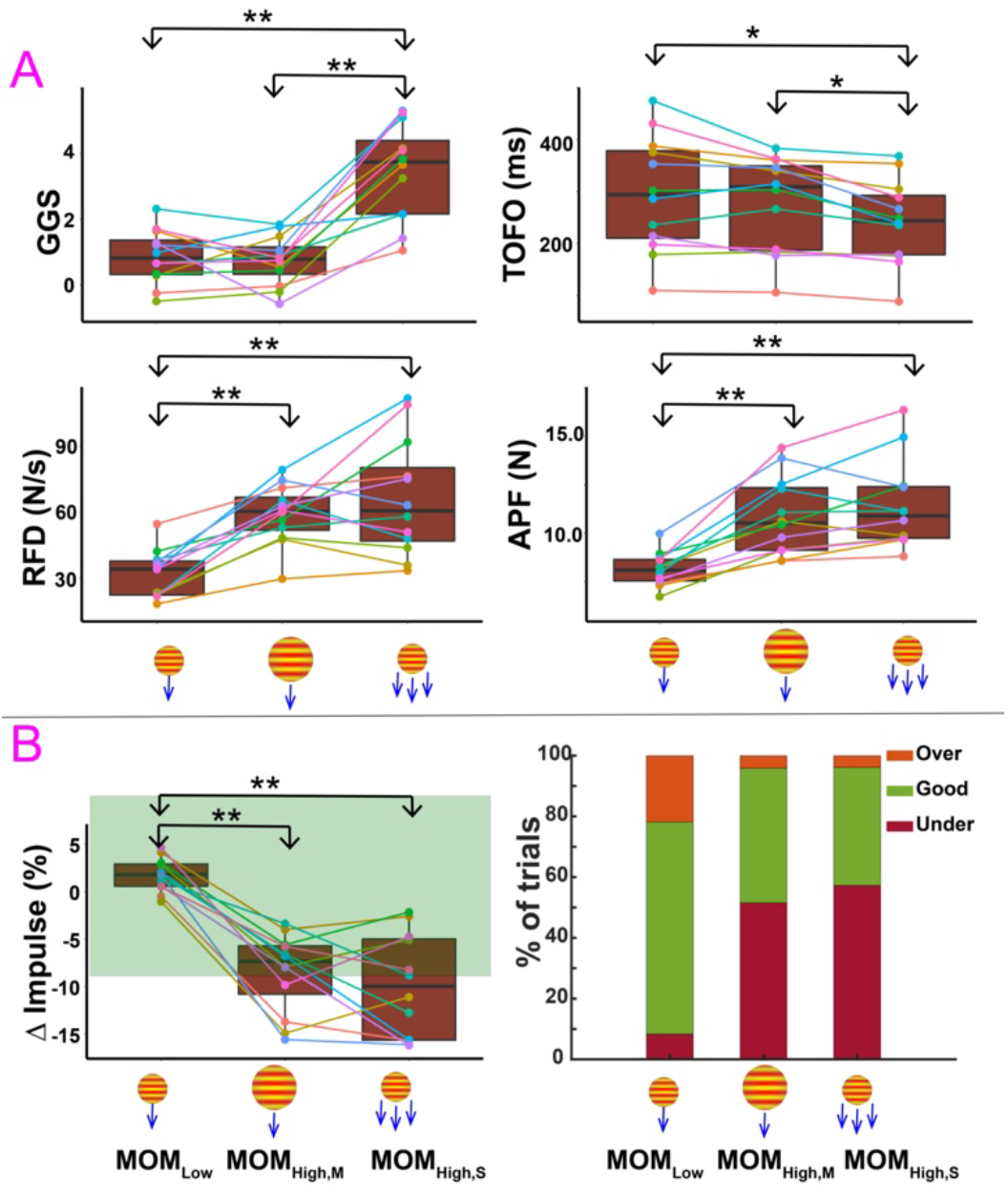
Target speed affected the timing of the motor response while target momentum affected anticipatory postural responses in Exp. 1. A) top left panels shows that the gaze gain slope (GGS) increased when speed of the target increased MOM_High,S_, but there were no changes when target momentum was increased by increasing the mass MOM_High,M_. The top right panel shows that the time of force onset (TOFO) decreased when target speed was higher (MOM_High,S_), but there was no change in TOFO when the virtual mass of the target was increased (MOM_High,M_). The bottom two panels shows that anticipatory postural responses, rate of force development (RFD) (left), and anticipatory peak force (APF) (right) increased regardless of how the target momentum was increased. B) The left panel shows the measure of task performance ΔImpulse (%). It decreased with an increase in momentum for both (MOM_High,M_, MOM_High,S_). The green region indicates the 8% error margin allowed in the study for ΔImpulse. The right panel shows that participants generally undershot the required impulse in both the high momentum conditions.

ΔImpulse decreased in magnitude at higher object momenta for both mass (MOM_High,M_) and speed (MOM_High,S_) (Fig. 5B) [main effect of momentum for ΔImpulse: *F*(2,22) = 50.54, *P* < 0.001, η^2^ = 0.64]. Post-hoc comparisons revealed a significant difference between MOM_Low_ and MOM_High,M_ and MOM_Low_ and MOM_High,S_ (p<0.01). During the higher momentum conditions, the participants more often undershot the needed force leading to a lower success rate.

### Constraining smooth pursuit eye movements (SPEM) in Exp. 2 only impacted anticipatory postural responses

Our results from Exp.1 suggest that SPEM may modulate the timing of the motor response (TOFO). This is consistent with the first prediction from our first hypothesis (H1). To test this hypothesis further, in Exp. 2, we constrained the gaze in certain blocks (shown as CONST blocks in Fig. 6). Eye movements were constrained by creating a designated area in which participants were instructed to fixate during the entire trial (see Fig. 1D). The average pursuit duration for the constrained gaze blocks (10.4±1.4%) was significantly less than the free-viewing blocks FV1 (83.4±2.4%) and FV2 (79.7±3.45%) (see Fig. 6A). We expected that when gaze was constrained, TOFO, RDF, APF, and ΔImpulse would all be affected because the limb motor system won’t have access to the extraretinal signals associated with SPEM.

RFD and APF decreased significantly (Figs. 6C and 6D). For RFD, there was a main effect of gaze condition (*F* (2,22) = 12.39, *P* < 0.001, η^2^ = 0.3). There was also a main effect of gaze condition for APF (*F* (2,22) = 20.39, *P* < 0.001, η^2^ = 0.33). The post-hoc tests revealed significant differences between the free-viewing (FV) and constrained gaze conditions for both RFD and APF (p<0.05). In contrast, TOFO showed no differences between the free-viewing and constrained-viewing conditions (Fig. 6B) [main effect of gaze condition for TOFO: *F* (2,22) = 0.39, *p* =0.68, η^2^ = 0.01]. In addition, compared to MOM_Low_ and MOM_High,M_, TOFO was reduced for MOM_High,S_ for both free-viewing and constrained conditions. This suggested that smooth pursuit eye movements (SPEM) perhaps may not play an essential role in modulation of the timing of the motor response (TOFO).

**Figure 6:**
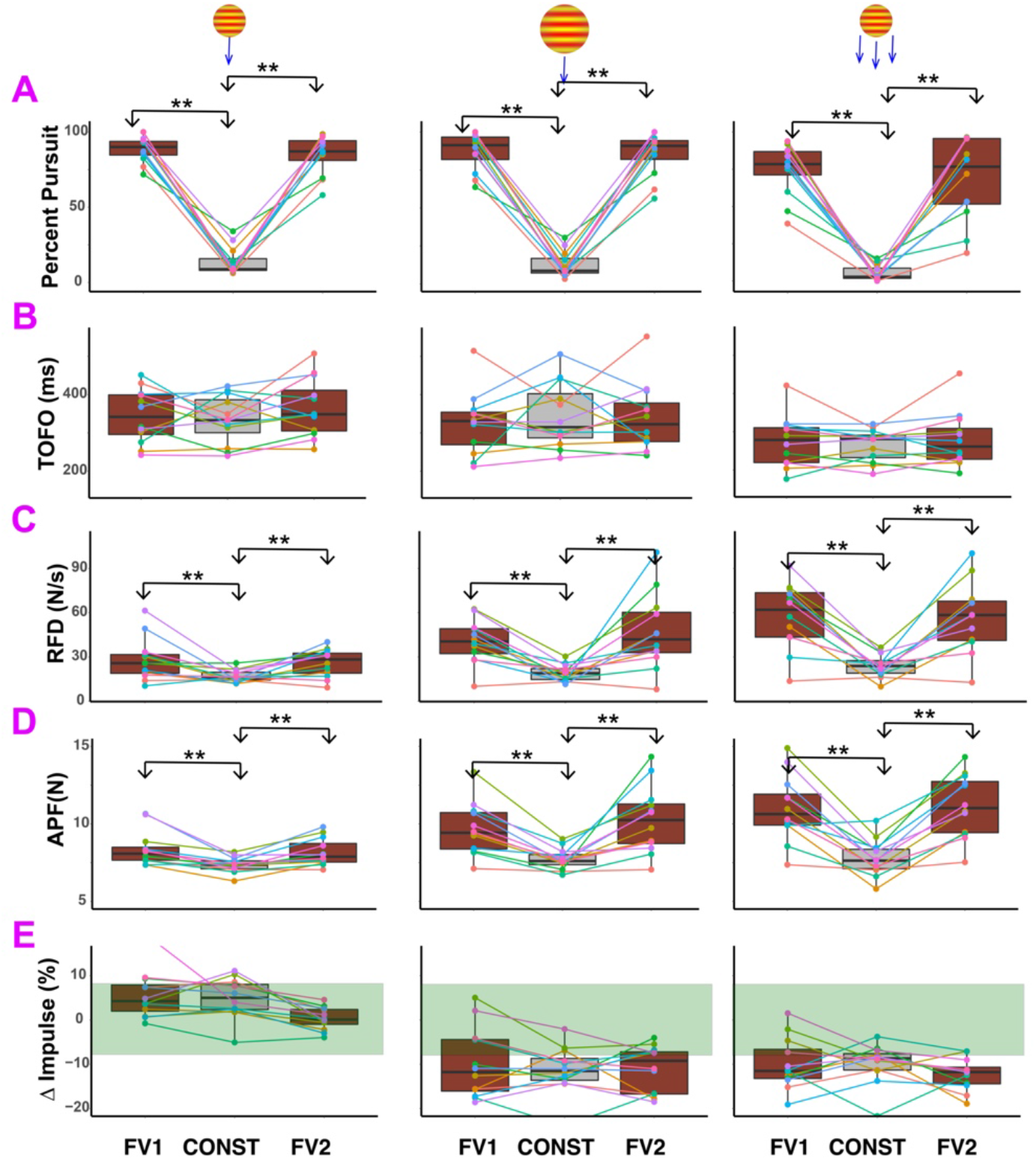
Constraining gaze selectively affected anticipatory postural responses. A) When gaze was constrained, the pursuit percentage (PP) dropped significantly for all three momentum conditions. B) Time of force onset (TOFO) showed no significant change when gaze was constrained. Important to note though is that TOFO was smaller for MOM_High,S_ compared to MOM_Low_ and MOM_High,M_ for both free-viewing and constrained conditions. This result is consistent with Exp.1. C and D) Both RFD and APF deceased significantly when gaze was constrained. E) ΔImpulse (%) did not show any modulation with constraining eye movements. But participants undershot ΔImpulse for the two high momentum conditions. This result is also consistent with Exp. 1. FV1 is first four blocks of free-viewing and FV2 is the last four blocks. CONST is the middle 8 blocks where gaze was constrained.

Finally, ΔImpulse was not impacted by constraining gaze (Fig 6E). There was no main effect of gaze condition on ΔImpulse (*F* (2,22) = 3.07, p*=*0.07, η^2^ = 0.06). Similar to Exp. 1, participants undershot the force required for performing the task correctly in the high momentum conditions of Exp. 2 (MOM_High,M_ and MOM_High,S_).

### Decreasing the reactive forces during contact in Exp. 3 affected motor response timing, anticipatory postural responses, and task performance

We compared the variables of interest between Exps. 2 and 3 using a one-way ANOVA with *force* as a factor (high and low as levels). For the same momentum/impulse combinations, in Exp. 3, the force applied by the robot was lower compared to Exp. 2 (see Fig. 1D) resulting in smaller reactive forces (but the force impulse and the object speed were the same as Exp. 2). We also confirmed the differences using the two-sample Kolmogorov-Smirnov (kstest) test.

Gaze gain slope (GGS) was unaffected by lowering the force in Exp.3 (Fig. 7A). There was no main effect of force for any of the momentum conditions. The kstest also showed that GGS in experiments 2 (high force) and 3 (low force) were not from different distributions. The one way-ANOVA for TOFO (see Fig. 7B) showed no main effect of force for MOM_Low_ (*F* (1,46) = 3.21, p=0.08). However, the main effect of force was significant for MOM_High,M_ (*F* (1,46) = 14.4, p*=* <0.001) and MOM_High,S_ (*F* (1,46) = 4.7, p<0.05). In contrast, the kstest showed that TOFO for all three momentum conditions emerged from different continuous distributions in experiments 2 (high force) and 3 (low force). This suggests that on average, the motor response is initiated earlier when the reactive forces generated during contact are smaller.

**Figure 7:**
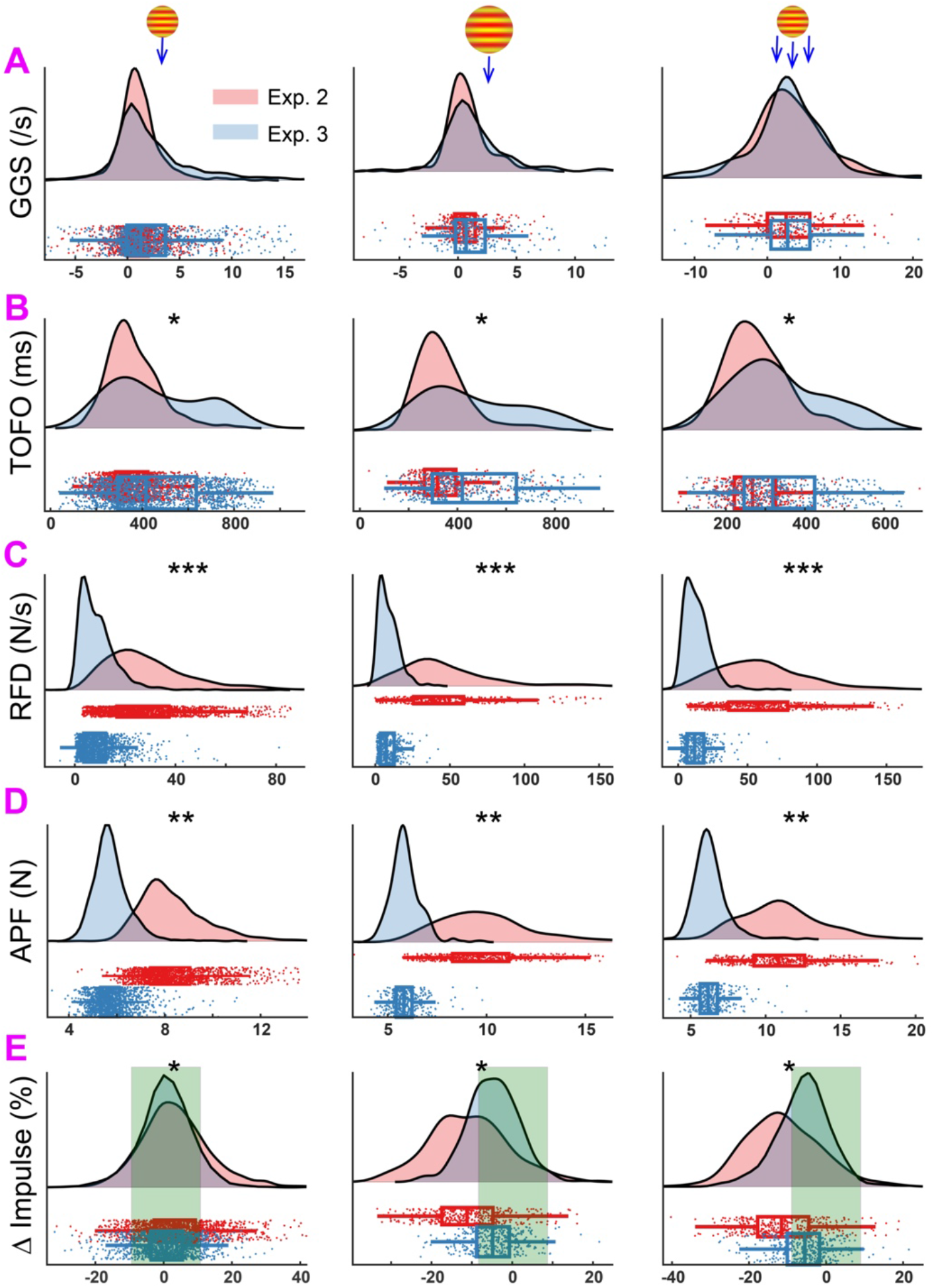
Softer collisions (lower reactive forces generated during contact) in Exp. 3 had upstream effects on the timing of the motor response and anticipatory postural variables, but not SPEM. The contact duration in Exp. 2 was 90 ms and in Exp. 3 was 160 ms. We performed both parametric and non-parametric tests because all assumptions for ANOVA were not satisfied. A) Gaze gain slope (GGS) did not show any significant difference between Exps. 2 & 3. B) TOFO was higher for Exp. 3. In other words, people increased hand force above baseline levels sooner when the collision was softer. C & D) Both the anticipatory posture variables, rate of force development (RFD) and anticipatory peak force (APF) decreases in Exp. 3. E) Participants were more accurate in Exp.3 than Exp. 2. ΔImpulse decreased and was closer to 0 in Exp. 3. The shaded green area indicates the 8% allowed error margin in our task.

The anticipatory response variables (RFD and APF) were significantly different between experiments 2 (high force) and 3 (low force). This was confirmed by the one-way ANOVA. The main effect of force was significant for RFD (see Fig. 7C) for all three momentum conditions [MOM_Low_ (*F* (1,46) = 76.36, p*=* <0.001), MOM_High,M_ (*F* (1,46) = 100.3, p*=* <0.001), & MOM_High,S_ (*F* (1,46) = 98.6, p<0.001)]. Similarly, the main effect of force was significant for APF (see Fig. 7D) [MOM_Low_ (*F* (1,46) = 152.8, p*=* <0.01), MOM_High,M_ (*F* (1,46) = 150.44, p*=* <0.01), & MOM_High,S_ (*F* (1,46) = 159.4, p<0.01)]. These differences between experiments 2 (high force) and 3 (low force) were also confirmed by the kstest. Finally, the one way ANOVA for ΔImpulse (see Fig. 7E) showed no effect of force on MOM_Low_ (p=0.07), but a main effect of force for both MOM_High,M_ (*F* (1,46) = 13.5, p*=* <0.001) and MOM_High,S_ (*F* (1,46) = 14.7, p<0.05) momentum conditions. These differences were also confirmed by the kstest.

### Peak hand reactive force during contact was a better predictor of overall task performance than anticipatory posture variables

The relationship between APF and ΔImpulse was not very strong (Fig. 4D), but APF scaled up for the high momentum conditions (MOM_High,M_ and MOM_High,S_) in both Exps. 1 (Fig. 5A) and 2 (Fig. 6D).

Furthermore, when reactive forces were lowered in Exp. 3, while keeping object speed and force impulse constant, both APF (Fig. 7D) and ΔImpulse (Fig. 7E) decreased. This suggested that perhaps the main role of APF was to stabilize posture against the large reactive forces generated during contact, but APF alone may not directly contribute to task performance.

To test this further, we performed a linear regression with ΔImpulse as the dependent variable and APF and peak hand reactive force (CPF) as predictor variables for both Exps. 2 and 3. We bootstrapped on the regression coefficients (see Methods) and found that the coefficients for APF were statistically indistinguishable from ‘0’, whereas the 95% confidence interval for coefficients of CPF did not include 0 (see Fig. 8). The regression model suggested that CPF was a predictor of ΔImpulse. We previously saw that APF and ΔImpulse were weakly correlated (Fig. 4D). APF was likely eliminated from the regression model because APF and CPF were also both correlated (data not reported), but CPF was likely a much stronger predictor of ΔImpulse.

**Figure 8:**
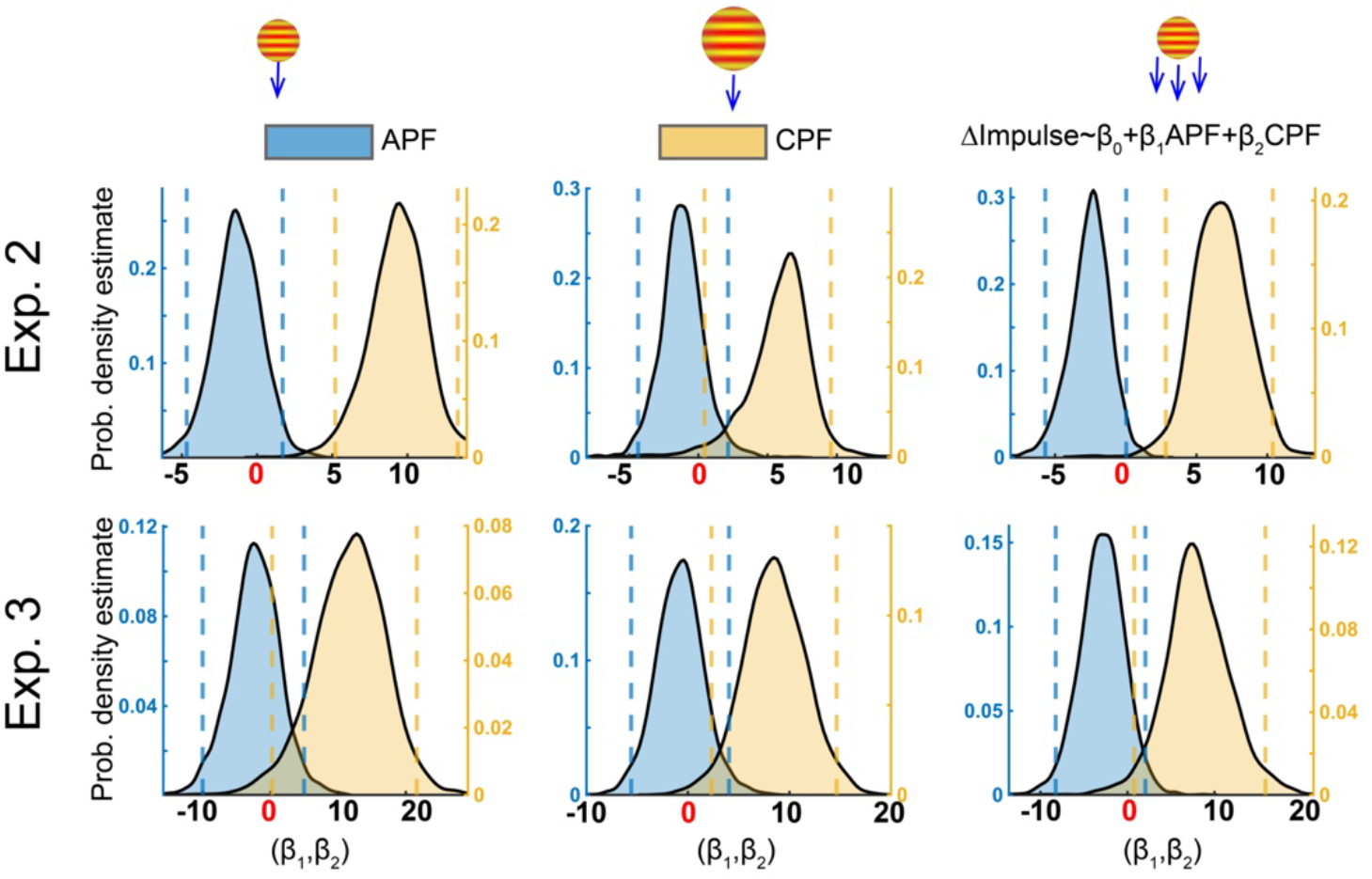
Peak reactive hand force during contact (CPF) is a better predictor of overall motor task performance than anticipatory posture variables peak force (APF). The plot of bootstrapped regression coefficients from the simple linear regression performed with ΔImpulse as the dependent variable and anticipatory peak force (APF) and the reactive peak force during contact (CPF) as predictor variables for both Exps. 2 and 3. The non-parametric confidence interval (CI) for the bootstrapped regression coefficients contains 0 for all the APF distributions (blue curve, both experiments all conditions) but the CI did not contain 0 for the CPF distributions (yellow curve, both experiments all conditions). The overlap with ‘0’ for APF suggests that this variable was not a significant predictor of ΔImpulse. This suggests that CPF is a stronger predictor of ΔImpulse than APF.

## Discussion

In this study, we conducted three experiments to probe the role of smooth pursuit eye movements (SPEM) on timing of motor responses (TOFO), anticipatory posture responses (RFD & APF), and task performance (ΔImpulse) in a task where participants had to stop a virtual moving object through modulation of reactive hand forces during contact between the object and the hand. We programmed the physics of mechanical interaction between moving objects and the body by assigning a mass to the virtual object and then by transforming the momentum (mass x speed) into a force-time template that the robot applied to the hand during the contact (31, 32). Participants were asked to stop the object by applying a force equal to the one applied by the robot during the contact. We call this paradigm the Mechanical Stopping of Virtual Projectiles (MSTOP). We proposed a hand force control model (Fig. 2) which articulated a hypothetical relationship between SPEM, timing of motor responses, anticipatory postural responses, and task performance. Our initial model was supported by the first experiment, but the subsequent experiments did not support this model.

### Participants learned the task not by decreasing the mean amplitude of ΔImpulse, but by decreasing the variability of ΔImpulse

Catching actions is an example of a complex motor skill that involves mechanical stopping of objects. These actions have both a kinematic and kinetic component. The kinematic component involves moving the arm and hand to an appropriate location in space to contact the object, and the kinetic component involves preparation of an anticipatory postural response to absorb the momentum of the ball during contact. The contact duration is typically too short for feedback and voluntary processes to have a meaningful role in modulation of reactive forces. The kinematic aspects of this skill have been well studied using interception paradigms (35, 48–51). Many studies, including from our group (35, 36) have used response latencies to define the features of motor planning and execution. Here, we focused primarily on the kinetic aspects of control.

The kinetic aspects of mechanical stopping of objects have been studied using catching experiments (17, 52, 53). These studies have revealed that a key feature of stopping objects is to prepare a feedforward motor response to stabilize hand posture in anticipation of contact between the object and the hand. These responses have been called anticipatory postural responses and studies have shown that these responses scale with object momentum (17). However, these studies did not measure the reactive interaction forces produced by the object and the hand during the contact. Using a virtual version of the task and a robotic manipulandum makes it possible for us to measure that and probe how this transient reactive force arises and how participants learn to modulate it with practice. ΔImpulse provides a metric of this interaction between the virtual object and the hand.

Our results suggest that participants improved performance not by minimizing the mean ΔImpulse, but by minimizing the variability of ΔImpulse across blocks (Fig. 3). This suggests that participants learned the task through motor exploration and as they learned the mechanics of the contact, the trial-by-trial and block-by-block variability decreased. Other studies have suggested that motor variability may be a fundamental mechanism of how the sensorimotor system learns new novel task dynamics (54, 55).

This view is based on reinforcement learning theory and equates motor variability with goal-directed and intentional exploration of the solution space. The exploration coupled with reinforcement could drive motor learning in MSTOP.

### Constraining smooth pursuit eye movements (SPEM) reduced the amplitude of anticipatory postural responses

Mechanical interactions between inertial objects and hand produce reactive forces. In catching studies, participants stabilize posture prior to contact between the object and the arm by co-contracting antagonist muscles (17, 56). In MSTOP, reactive forces are produced by the robot on the hand and vice-versa. Participants stabilized hand posture by increasing hand force prior to contact. Important to note is that like catching actions, the anticipatory forces have no direct impact on task performance because these forces are produced prior to the contact. Thus, the anticipatory increase in hand force only stabilizes posture against the large reactive forces generated during contact.

The reactive force produced by the hand is largely determined by the force applied by the robot – the larger the robot force, the larger the reactive force produced by the hand. The short duration of the contact in Exps. 1 & 2 (90 ms) makes it very difficult to voluntarily modulate this force during the contact. Typically fast feedback corrections are observed in muscle activity within the first 50-100 ms after the onset of a mechanical perturbation (57–59), but hand force is likely low-pass filtered by the musculoskeletal system (60) and consequently may not include effects of feedback responses. It is extremely unlikely that the short time window of contact in Exps. 1 & 2 allowed voluntary modulation of force. Thus, the hand force during contact may be largely determined by feedforward control and intrinsic viscoelastic properties of muscles.

Our overall proposal was that anticipatory posture variables (RFD and APF) and the time at which hand force was increased in anticipation of the contact (TOFO) were modulated by smooth pursuit eye movements (SPEM). Specifically, our first hypothesis (H1) was that SPEM contribute to adaptive modulation of TOFO, RFD, APF, and ΔImpulse. The two predictions from our first hypothesis were: 1) in free-viewing conditions, higher SPEM speeds to track faster moving objects would increase TOFO, but RFD, APF, and ΔImpulse would increase regardless of whether the object momentum increased because of mass or speed; and 2) constraining eye movements would decrease TOFO and increase RFD, APF, and ΔImpulse. We proposed the hand force control model (Fig. 2) to predict how SPEM would affect TOFO, RFD, APF, and ΔImpulse.

Trial-by-trial correlations in Exp. 1 between pairs of variables supported the hand force control model (Fig. 4). We also manipulated object momentum in Exp. 1 by either increasing its mass (MOM_Low_ to MOM_High,M_) or speed (MOM_Low_ to MOM_High,S_). Contrary to our first prediction, we found that increasing the speed of the object caused an increase in the slope of the gaze gain (Fig. 5) and caused the motor response to be initiated closer to contact between the object and the hand (decreased TOFO).

Consistent with our prediction, RFD and APF both increased in response to object momentum regardless of whether the momentum increased because of mass (M_High,M_) or speed (M_High,S_). Finally, ΔImpulse increased in magnitude but became more negative when the object momentum was increased.

The moderate to strong correlations that were observed between pairs of variables in Exp.1 for the hand force control model were necessary, but not sufficient for establishing stronger causal relationships between these variables. To that end, we constrained eye movements in Exp. 2 (Fig. 6). Object movement in a world-centered reference frame requires that the visual system integrate motion signals derived from retinal slip with an efferent copy of extraretinal ocular motor signals (1, 61–63). We expected that if gaze movements are constrained and the extraretinal contribution is suppressed, then the retinal slip of the object alone may cause over-estimation of object speed (22, 23). This should have increased the amplitude of the anticipatory postural responses (RFD and APF). Our results were not consistent with this prediction. We found that RFD and APF amplitudes decreased when SPEM were constrained for all three momentum conditions. This supports the idea that the hand motor system may have access to efferent copies of ocular motor signals and if those signals are suppressed, then the hand motor system underestimates the anticipatory postural responses needed to stabilize the hand against reactive forces. Surprisingly, the changes in anticipatory postural responses during constrained viewing did not affect task performance (ΔImpulse). These results suggest that anticipatory modulation of posture may be critical for stabilizing posture against reactive forces but may not assist with task performance. Furthermore, these results suggested that anticipatory postural responses and “compensatory” processes during the contact period may perhaps be independently modulated by the nervous system.

### Gaze gain signals may have a minimal effect on hand motor control

One of the interesting findings was that gaze gain was modulated with different object speeds. A gaze gain of unity at every time point would ensure that the gaze remains on the moving object. But since the objects were moving towards the body and we know that SPEM are slower when combined with vergence eye movements (64, 65), we expected gaze gain may change during the trial. Thus, we calculated gaze gain slope (GGS) instead of gaze gains. Theoretically, there was no reason for this slope to differ between the three momentum conditions, but the gaze gain slope (GGS) was higher for MOM_High, S._ This suggested that the faster object speed in the MOM_High,S_ condition placed greater demands on the ocular motor system. After Exp. 1, we believed that this change in GGS for higher object speeds facilitated a late hand motor response (shorter TOFO), but after Exps. 2 & 3, where we observed that TOFO was not affected by constraining gaze, we discarded this possibility. In fact, it appears that the gaze gain signal may not be contributing to hand force control. The only SPEM signal that mattered for hand force control was whether the gaze pursued the object -if the gaze was constrained, anticipatory postural responses were subdued (smaller in magnitude).

SPEM gains have been reported in hundreds of studies and are considered to be an important neurophysiological biomarker for understanding the limits of the ocular motor system itself and how specific neurological (66) and psychiatric diseases (67) affect this system. While it is known that manual tracking of moving objects increases the gain of SPEM (68, 69), it is unclear if higher SPEM gains also increase movement accuracy. One reason for that is that SPEM gains are relatively preserved for an object moving at the same speed across trials. Therefore, it is unclear how the gain signals might contribute to parameters of hand force control. In contrast, studies that use occlusion or where gaze is constrained show that limb motor performance is much more accurate when participants are allowed to pursue moving objects (20, 25). Together, this suggests that perhaps the pursuit signal itself may be sufficient for improving hand motor control during interactions with moving objects, but the pursuit or gaze gain signal may not be critical for precise control of hand force.

### Independent modulation of APFs and ΔImpulse suggests a switch in control policy during contact

When participants catch balls falling freely, they exhibit a unique kinematic signature -wrist flexors are activated right before contact between the object and the hand. Participants flex the wrist to initiate hand movement towards the object approximately 100-150 ms before contact with the object (17, 56, 70). This anticipatory flexion of the wrist seems to be driven primarily by visual processing of moving objects (71). We observed similar kinematics in our study (data not presented) where every participant moved the hand towards the approaching object by increasing hand force in the Y-direction starting at TOFO. It is unclear why participants in our study and the abovementioned studies moved their hand towards the approaching object, but many participants in our study reported that they found that they performed better if they moved their hand towards the object.

There is no mechanical necessity to move the hand towards the object – participants could stop/catch the objects by simply keeping the hand steady. However, it appears that the initial movement towards the object facilitates modulation of reactive forces during contact. A study done by Cesari, Bertucco, and colleagues has shown that when participants stop a pendulum during upright stance, in a 200 ms window before contact, they reciprocally activate antagonist pairs of postural muscles (72), i.e., one muscle is activated while its antagonist is inhibited. They also showed that during impact [50 to 300 ms after contact], the motor system switched to a co-activation mode where antagonist pairs were simultaneously co-activated. The switch in control policy may be critical for simultaneous posture stabilization and task success and may be the reason why APF and ΔImpulse did not exhibit parallel modulation in Exps. 2 & 3 (Fig. 6D & E). Venkadesan and colleagues have used electromyographic signals to show switches in control policies when humans move their fingers from motion to static force production (73). Our task was similar in spirit in the sense that participants moved their hands right before contact between the object and the hand. This switch in control policy may be mediated by the premotor cortex (74).

The relative independence of APF and ΔImpulse is also supported by comparing how the anticipatory peak force (APF) and the contact peak force (CPF) affected ΔImpulse in Exps. 2 and 3 using a regression analysis (Fig. 8). We made the collision “softer” in Exp. 3 by reducing the reactive force generated by the hand during contact while keeping the object speed and the force impulse applied by the robot during the contact the same. The model showed that CPF is a much stronger predictor of ΔImpulse, though ΔImpulse and APF were also correlated. Furthermore, we found that RFD and APF were lower and ΔImpulse was smaller in Exp. 3 (Fig. 7). Together, these results show anticipatory force responses scale according to the magnitude of the contact forces – the larger the contact force, the larger the anticipatory force. These results also raise the possibility that needs to be tested further – anticipatory postural responses scale based on the magnitude of the contact forces but may not be necessary for accurate task performance.

### Extraretinal pursuit signals may contribute to modulation of anticipatory postural responses, while retinal motion signals may be more important for timing of motor response

Based on the results of the three experiments, we modified our original hand force control model (Fig. 2). This new model is shown in Fig. 9. The new model incorporates key results from all three experiments. First, we observed that constraining the gaze movements in Exp. 2 had no effect on the timing of the motor response (TOFO, Fig. 6), but TOFO decreased when objects moved faster in both Exps. 1 & 2 (MOM_High,S_ condition, Figs. 5 & 6), regardless of whether the gaze was constrained or not. A recent study has shown that 300 ms of viewing time is sufficient to initiate a timely motor response (75). Together, these results suggest that extraretinal signals associated with SPEM may not be necessary for the precise timing of motor responses; retinal signals of object motion alone may be sufficient.

**Figure 9:**
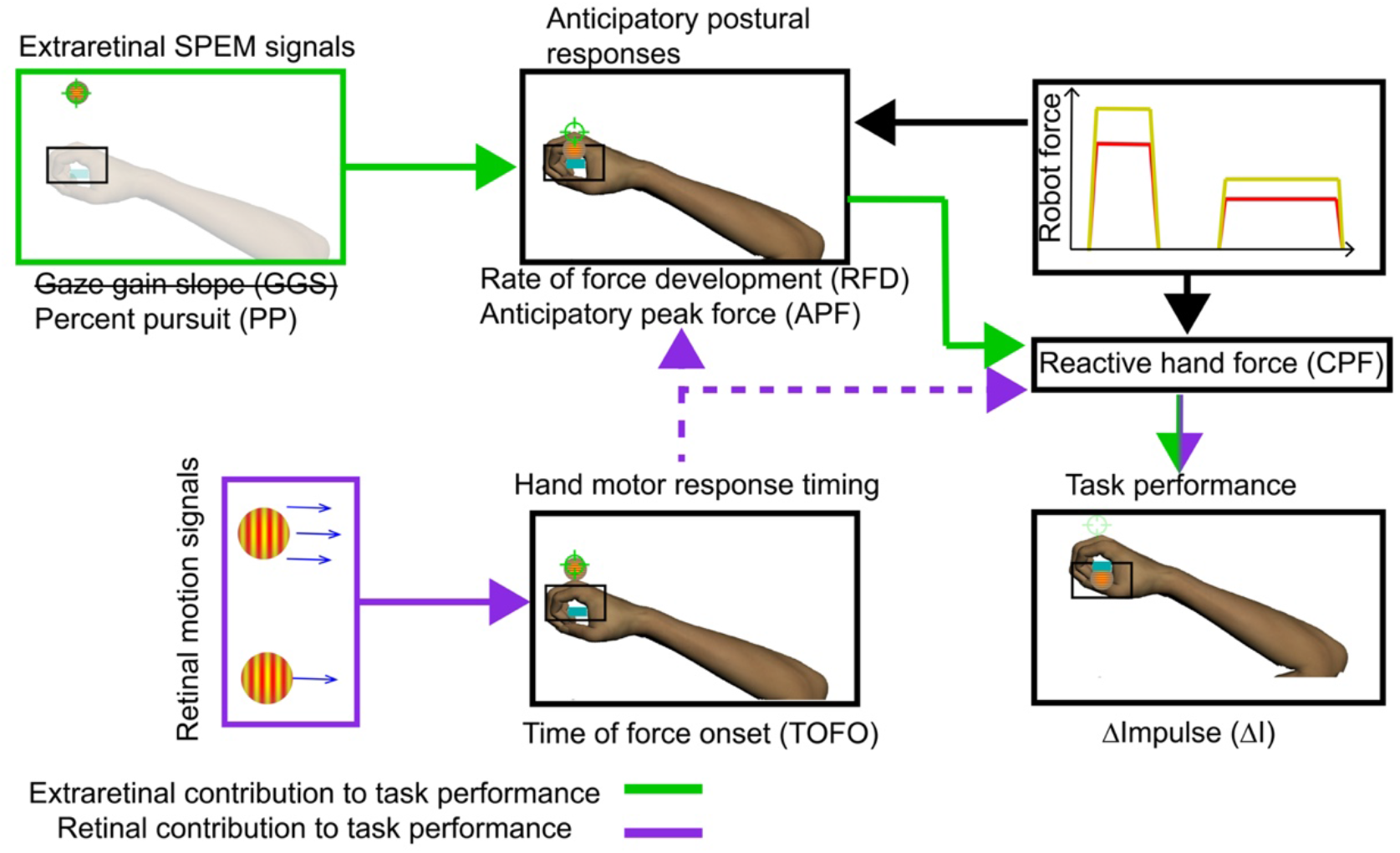
Modified hand force control model. The extraretinal signals associated with smooth pursuit eye movements (SPEM) seemed to only influence anticipatory postural variables (solid green arrow). The anticipatory postural variables themselves were weakly correlated with ΔImpulse, but anticipatory postural variables decreased in magnitude when gaze was constrained while ΔImpulse did not. This suggested that anticipatory posture stabilization may indirectly affect task performance though modulation of reactive hand force (CPF). The time of force onset (TOFO) was sensitive to target speed regardless of whether participants pursued the targets or when their gaze was constrained. This suggested that retinal motion signals may be sufficient to drive the timing of the motor response (solid violet arrow). The correlations between TOFO and RFD also suggest that the timing of the motor response may have an indirect effect on posture stabilization. Retinal motion signals may contribute to reactive forces produced by the hand through this mechanism (dashed violet arrows). This needs to be probed further.

Constraining gaze affected anticipatory postural responses. Specifically, when gaze was constrained, the rate of force development (RFD) and anticipatory postural force (APF) were lower (Fig. 6), suggesting that the anticipatory postural responses were weaker in the absence of the extraretinal input. Surprisingly, the weaker responses had little to no effect on task performance, i.e., ΔImpulse was unaffected in the absence of SPEM. This strongly implicates APF with posture stabilization and not task performance.

The third experiment (Exp. 3) helped us better understand why anticipatory postural responses had negligible effects on task performance. Here, we lowered the reactive forces during contact while keeping object motion and overall force impulse the same. We hypothesized that upstream gaze and hand motor variables would be modulated by this change (H2). We found that APF was lower in Exp. 3 and task performance was more accurate (lower ΔImpulse). TOFO also increased but gaze gains remained unchanged in Exp. 3. These results partially supported H2. Together, this suggests that extraretinal signals may be important for posture stabilization but perhaps not task performance, whereas retinal motion signals may indirectly affect task performance by modulating the timing of the motor response and hand force. This needs to be tested further.

### The cerebellum may modulate anticipatory postural responses through SPEM

The cerebellum is a critical node for the control of smooth pursuit eye movements (SPEM) (76, 77). Extrastriatal and frontal visual areas project to the pontine nuclei and send both ocular motor and visual information signals. That information is then sent from the pontine nuclei to the paraflocculus, the flocculus, and the vermis in the cerebellum. The processed output of the cerebellum is projected back via the deep cerebellar nuclei to the vestibular nuclei and the oculomotor nuclei for control of SPEM. There are also strong reciprocal connections between the cerebellum and the cortical areas, specifically the frontal eye fields, that play a pivotal role in the control of SPEM. Damage to the cerebellum is known to cause deficits in SPEM (78–81).

In a study performed by Lang and Bastian, cerebellar patients were asked to catch balls of different weights dropped from a fixed height. The patients were unable to increase the anticipatory muscle activity in preparation of contact between the ball and the hand (56, 82). In contrast, healthy controls were easily able to modulate anticipatory muscle activity in the flexor muscles of the arm to absorb the ball momentum at contact. Lang and Bastian also showed that the cerebellar patients were unable to modulate anticipatory muscle activity to balls of different weights. This led the authors to conclude that the cerebellum is a critical node for appropriate tuning of anticipatory responses to task demands. This conclusion is supported by an imaging study that showed that the cerebellum is a key neural substrate involved in trial-by-trial modulation of feedforward motor commands for catching actions (83). The results from these studies and our own raise the possibility that in tasks that require interactions with moving objects (e.g., catching), the cerebellum may play a critical role in anticipatory posture stabilization by processing extraretinal motion signals associated with SPEM.

In conclusion, our results demonstrate that extraretinal signals associated with SPEM may play an important role in anticipatory stabilization of posture prior to mechanical interactions with moving objects, whereas retinal motion signals may play a stronger role in modulating the timing of the motor responses. We also found that it was not gaze gain (ratio of gaze and object speed) signal that impacted posture stabilization, but whether the gaze itself was engaged in smooth pursuit of objects. Surprisingly, we also observed that the magnitude of reactive forces that had to be produced by participants during contact had a strong impact on the timing of motor responses and anticipatory modulation of posture. SPEM and anticipatory stabilization of hand posture did not appear to have a significant impact on task performance. This suggests that the old adage “keep your eyes on the ball” may be more important for stabilizing posture during mechanical interactions with moving objects rather than the task performance itself.

